# Feeding Neurons Integrate Metabolic and Reproductive States in Mice

**DOI:** 10.1101/2023.01.25.525595

**Authors:** Megan G. Massa, Rachel L. Scott, Alexandra L. Cara, Laura R. Cortes, Norma P. Sandoval, Jae W. Park, Sahara Ali, Leandro M. Velez, Bethlehem Tesfaye, Karen Reue, J. Edward van Veen, Marcus Seldin, Stephanie M. Correa

## Abstract

Trade-offs between metabolic and reproductive processes are important for survival, particularly in mammals that gestate their young. Puberty and reproduction, as energetically taxing life stages, are often gated by metabolic availability in animals with ovaries. How the nervous system coordinates these trade-offs is an active area of study. We identify somatostatin neurons of the tuberal nucleus (TN^SST^) as a node of the feeding circuit that alters feeding in a manner sensitive to metabolic and reproductive states in mice. Whereas chemogenetic activation of TN^SST^ neurons increased food intake across sexes, selective ablation decreased food intake only in female mice during proestrus. Interestingly, this ablation effect was only apparent in animals with a low body mass. Fat transplantation and bioinformatics analysis of TN^SST^ neuronal transcriptomes revealed white adipose as a key modulator of the effects of TN^SST^ neurons on food intake. Together, these studies point to a mechanism whereby TN^SST^ hypothalamic neurons modulate feeding by responding to varying levels of circulating estrogens differentially based on energy stores. This research provides insight into how neural circuits integrate reproductive and metabolic signals, and illustrates how gonadal steroid modulation of neuronal circuits can be context-dependent and gated by metabolic status.

## Introduction

The homeostatic processes of metabolism and reproduction are mutually dependent on one another. Reproductive milestones, including pregnancy and quiescence, result in major metabolic shifts (Baz et al., 2016; Mauvais-Jarvis, 2017; Palmer and Clegg, 2015), and metabolic status can gate reproductive function in menstrual and estrual mammals. Pubertal onset has been shown to require a critical threshold of body fat (Frisch, 1972; Frisch and Revelle, 1970), and increasing adiposity is associated with a decreasing age of pubertal onset in individuals with ovaries in particular (Solorzano and McCartney, 2010). During reproductive years, undernutrition can acutely disrupt menstrual cyclicity (Nelson et al., 1985) via hypogonadotropic hypogonadism (Muñoz and Argente, 2002) and result in fewer successful pregnancies (Ball et al., 1947). Rodent models have been used to investigate the effects of sex variables on metabolic tissues. Estradiol has been shown to regulate key metabolic processes such as adiposity (Palmer and Clegg, 2015), thermogenic capacity & locomotion (Correa et al., 2015; Martínez de Morentin et al., 2014; Musatov et al., 2007; Palmer and Clegg, 2015; Xu et al., 2011), and feeding (Massa and Correa, 2020). Sex chromosome complement has been demonstrated to affect fat deposition and energy output (Chen et al., 2012). Few studies have investigated the mechanisms by which both metabolic and reproductive status reciprocally interact to modulate behavior.

A candidate region for the seat of this interaction is the mediobasal hypothalamus, a region typically thought to be comprised of the arcuate nucleus, ventromedial nucleus, and median eminence. Positioned near the third ventricle and partially outside of the blood brain barrier (Ciofi, 2011), the mediobasal hypothalamus is situated as a nexus region with the ability to sample circulating homeostatic hormones and relay relevant information to other regions of the brain. Importantly, this region is sensitive to and vital for both reproductive and metabolic homeostasis. For instance, in addition to the role of kisspeptin neurons in the arcuate nucleus as key modulators of estrogen-mediated negative feedback in the hypothalamic-pituitary-ovarian axis (Mittelman-Smith et al., 2012; Smith et al., 2005), they also require epigenetic de-repression for menarche (Lomniczi and Ojeda, 2016; Wright et al., 2021). The arcuate nucleus is also home to the canonical homeostatic feeding neurons – appetitive agouti-related peptide (AgRP)/neuropeptide Y (NPY) neurons and satiety-related proopiomelanocortin (POMC) neurons – which can detect and respond to sensory and metabolic cues including leptin, ghrelin, and insulin (Burnett et al., 2016; Chen et al., 2015; Oldfield et al., 2016). The ventromedial nucleus of the hypothalamus, and in particular the ventrolateral subregion, is important for various functions across sexes, including mating behavior (reviewed in (Kammel and Correa, 2019)). Distinct, estrogen-sensitive cellular populations in the ventrolateral ventromedial nucleus contribute to various aspects of metabolism, including locomotion (Correa et al., 2015; Krause et al., 2021; Narita et al., 2016) and thermoregulation (van Veen & Kammel et al., 2020). Given the substantive neuronal heterogeneity within a small brain region, it is unsurprising that these populations form complex functional interactions. For example, simulation of starvation via chronic chemogenetic activation of appetitive arcuate AgRP neurons has been demonstrated to acutely disrupt estrous cyclicity (Padilla et al., 2017), providing a neuronal contributor to the link between metabolic state and reproductive function.

The tuberal nucleus (TN) straddles the mediobasal hypothalamus and lateral hypothalamic area. Itis an understudied region marked by expression of the neuropeptide somatostatin (SST) which has been mostly characterized in rats. Studies have found the TN to be closely related to the ventrolateral region of the ventromedial hypothalamus (Canteras et al., 1994) (though in a recent mouse study, the TN was considered a part of the lateral hypothalamic area; Mickelsen et al., 2019), and suggest the region may also be estrogen sensing (Canteras et al., 1994; Pfaff and Keiner, 1973; Simerly et al., 1990). Recent studies investigating this region in mice have found the TN to promote feeding behavior through canonical homeostatic feeding circuitry (Luo et al., 2018) and learning/hedonic feeding in males (Mohammad et al., 2021). However, this region is an excellent prospective candidate for integrating metabolic and reproductive cues to affect feeding based on its anatomical location, possible sensitivity to circulating reproductive hormones, detection of metabolic hormones such as ghrelin (Luo et al., 2018), and promotion of feeding behavior. Indeed, whole-body knockouts of SST exhibit weight gain that is exacerbated by sex category and high fat diet (Luque et al., 2016).

Here, we use mice to interrogate the role of SST neurons of the tuberal nucleus (TN^SST^) in integrating metabolic and reproductive cues to affect feeding. TN^SST^ neurons exhibited differential control of feeding in female and male mice (defined by anogenital distance at weaning and postmortem inspection of the gonads), with neuronal ablation decreasing food intake only in females. This effect was primarily due to a decrease in food intake during proestrus, when circulating ovarian hormones are at high concentrations, and was only observed in animals with a low body mass. To determine whether adiposity could mediate the effect of body mass on food intake, fat transplantation experiments were performed. Increased white adipose tissue was sufficient to induce TN^SST^ neuronal modulation of food intake. Together, these data reveal a context dependent role for TN^SST^ neurons in the regulation of food intake, by which TN^SST^ neurons tune feeding behavior in response to metabolic and reproductive states.

## Results

### Chemogenetic activation of TN^SST^ neurons increases food intake in female and male mice

To test the role of TN^SST^ neurons across sexes, an AAV expressing a Cre-dependent Gq-coupled hM3Dq (Krashes et al., 2011) was stereotaxically injected to the TN of *Sst-Cre* mice (Figure 1A-C). Overall, activation of TN^SST^ neurons using the small molecule ligand clozapine-N-oxide (CNO) increased daytime food intake over a four-hour testing period in both females and males (Figure 1D). Control animals without expression of hM3Dq-mCherry confirmed no effect of CNO alone, while within-subjects comparisons of animals expressing hM3Dq in TN^SST^ neurons indicated an increase in feeding upon CNO-induced cellular activation. There was a significant interaction between hM3Dq presence and treatment (saline v. CNO), both with time (treatment*hM3Dq*time, F(2,105)=3.2964, p=0.0409) and without time (treatment*hM3Dq, F(1,105)=35.2054, p<0.0001). The effect of neuronal activation (genotype-by-treatment interaction) was most prominent across sexes at 4 hours post-CNO injection (females: F(1,12)=10.0208, p=0.0081; males: F(1,9)=12.1521, p=0.0069), though males also exhibited a significant activation-by-treatment interaction at 2 hours post-CNO administration (F(1,9)=8.6957, p=0.0163). Post-hoc within-subjects analyses of animals bilaterally transduced with hM3Dq-mCherry indicated a significant increase in food intake during activation by CNO as compared to treatment with saline control (females overall: t(31)=2.8486, p=0.007732; males at 2 hours: t(5)=2.9701, p=0.0311; males at 4 hours: t(5)=3.3263, p=0.0286). Thus, activating TN^SST^ neurons elicits feeding across sexes.

**Figure 1:**
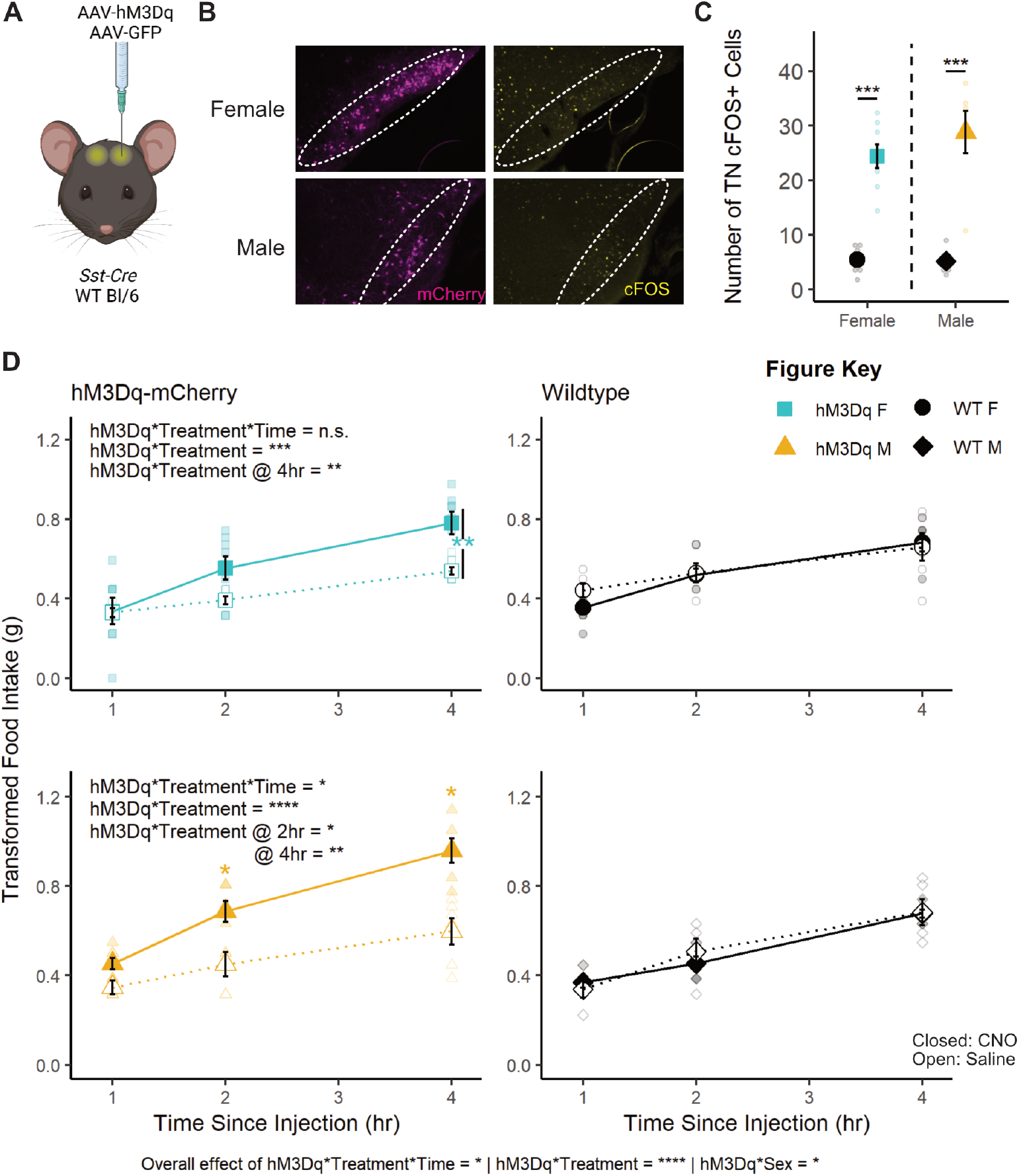
Transient activation of TN^SST^ neurons increased food intake across sexes. (A) Schematic of experimental paradigm. AAVs encoding hM3Dq-mCherry or GFP within Flip-Excision (FLEX) cassettes were injected bilaterally into the tuberal nucleus (TN) of *Sst-Cre* or wildtype mice. Created with BioRender.com. (B) Fluorescent images of mCherry (magenta) and cFOS (yellow) in the TN of *Sst-Cre* mice injected with h3MDq-mCherry show mCherry expression and more cFOS positive cells 90 minutes after CNO injection regardless of animal sex (overall effect of hM3Dq-mCherry presence, F(1,21)=73.634, p<0.0001; no interaction effect, F(1,21)=0.939, p=0.3435). Dotted line indicates boundaries of the TN. (C) Quantification of cFOS+ cells in the TN. Activation of TN^SST^ neurons in both female and male mice leads to higher food intake within the four-hour daytime testing period (left column, interaction between hM3Dq-mCherry presence and treatment: F(2,105)=3.2964, p=0.0409). (D) Within sex, only males exhibited an effect of activation and treatment over time (F(2,45)=3.2793, p=0.0468), though both females and males exhibited an effect of hM3Dq presence and treatment independent of time (F(1,60)=12.7928, p=0.0007 and F(1,45)=25.1794, p<0.0001, respectively). Males also exhibited a significant hM3Dq-by-treatment interaction at 2 hours post-CNO (F(1,9)=8.69, p=0.0163). CNO did not affect food intake in wild-type control mice (right column). Mean ± SEM, *p<0.05, **p<0.01, ***p<0.001, ****p<0.0001. F Control n=6; M Control n=5, F hM3Dq n=8; M hM3Dq n=6.

### Caspase ablation of TN^SST^ decreases food intake only in females

To determine if permanent inactivation of TN^SST^ alters feeding across sexes, an AAV expressing a Cre-dependent modified caspase virus (taCasp3-TEVp; (Yang et al., 2013) was stereotaxically delivered to the TN of *Sst-Cre* mice (Figure 2A). Bilateral elimination of *Sst* expression was validated by *in situ* hybridization (Figure 2B). Mice were subjected to two 96-hour food assays along with a battery of other metabolic tests. Final food intake, accounting for spillage, is depicted as an average over 24 hours (Figure 2C). ANOVA revealed an overall effect of sex where males consume more food than females, as expected (F(1,38)=14.1896, p=0.0006).

**Figure 2:**
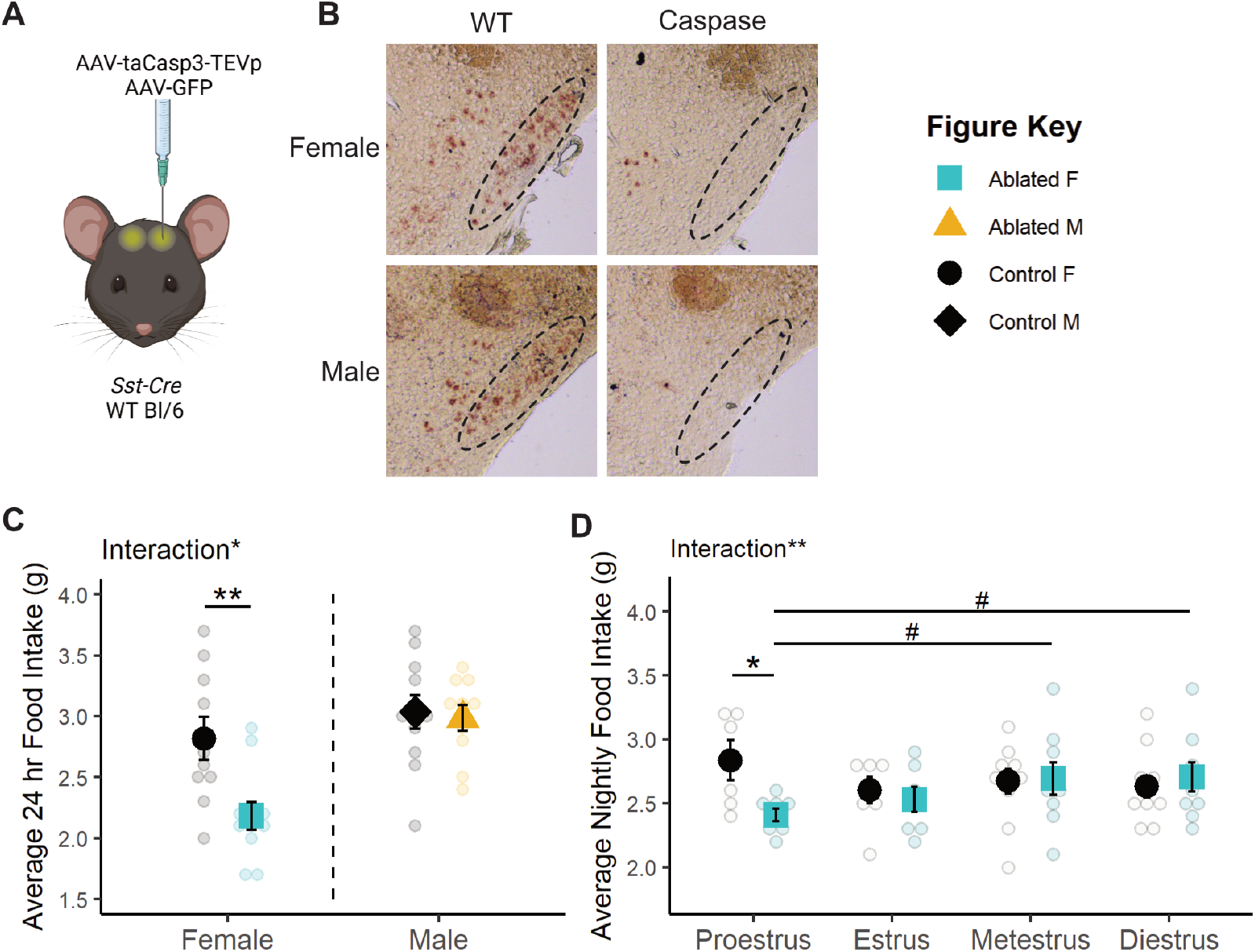
Caspase ablation of TN^SST^ neurons decreased food intake only in females. (A) Schematic of experimental paradigm. AAVs encoding taCasp3-TEVp or GFP within Flip-Excision (FLEX) cassettes were injected bilaterally into the tuberal nucleus (TN) of *Sst-Cre* or wildtype mice. Created with BioRender.com. (B) Representative brightfield images of *Sst* transcript expression in the TN of caspase-ablated and control animals. Dotted line indicates boundary of the TN. (C) Permanent TN^SST^ neuronal ablation decreases average daily food intake in females but not males. F Control n=10; M Control n=11; F Ablated n=11; M Ablated n=10. (D) This decrease in food intake is detected only in the night of proestrus. Proestrus: Control n=6, Ablated n=8; Estrus: Control n=7, Ablated n=7; Metestrus: Control n=10, Ablated n=9; Diestrus: Control n=10, Ablated n=9. Mean ± SEM; between subjects: *p<0.05, **p<0.01; within subjects: ^#^p<0.10.

Interestingly, the effect of TN^SST^ neuron ablation differed by sex (sex-by-ablation interaction: F(1,38)=4.6852, p=0.0368), an effect which post-hoc t-tests revealed to be carried by a decrease in food intake specifically in females (t(15.671)=−3.0561, p=0.007686, Figure 2C). This sex difference was not previously reported, but collapsing these data across sex results in an overall decrease in food intake with TN^SST^ neuron ablation (t(39.86)=−2.2713, p=0.0286), consistent with previous studies (Luo et al., 2018).

The female specific effect of TN^SST^ neuron ablation is modulated by estrous cycle stage. There was a significant interaction between TN^SST^ neuron ablation and estrous stage (F(1,45)=8.1581, p=0.0065) which was predominantly due to a decrease in food intake specifically during the night of proestrus (t(5.902)=−2.6044, p=0.04104, Figure 2D). Within-subjects analysis of mice in the neuronal ablation group suggested that nighttime food intake during proestrus was also slightly lower than consumption during metestrus (t(7)=−1.976, p=0.0887) and diestrus (t(7)=−2.3276, p=0.0528), although these effects across the cycle did not reach statistical significance.

Despite effects on food intake, no other metabolic measures were altered by TN^SST^ neuron ablation (see Table 2 for statistical results). TN^SST^ neuron ablation did not affect telemetry measures of activity/movement (Supp Fig 1A) or core body temperature (Supp Fig 1B), nor response to fasting glucose tolerance test (Supp Fig 1C). TN^SST^ neuron ablation did not affect body mass (Supp Fig 1D), suggesting that the selective decrease in food intake during proestrus was not sufficient to alter body mass.

### Body mass, specifically adiposity, influences effect of TN^SST^ neuronal ablation on food intake

There was an overall interaction between body mass and estrous phase (F(1,150)=3.9433, p=0.04889) and between body mass and ablation (F(1,150)=16.9924, p<0.0001; see Table 2 for full results). Analyzing the relationship between body mass and food intake within each estrous stage revealed significant negative correlations between body mass and food intake in wild-type animals during nights of proestrus (r^2^=0.3263; F(1,17)=8.235, p=0.01062; Figure 3A) and metestrus (r^2^=0.2801; F(1,16)=6.227, p=0.0239; Supp Fig 2), stages with higher relative circulating estradiol levels (Handelsman et al., 2020). There were no correlations between food intake and body mass in estrus and diestrus, stages with lower relative circulating estradiol levels (Figure 3A, Supp Fig 2, Table 2). TN^SST^ neuron ablation uncoupled this relationship with body mass in high estradiol stages (proestrus: r^2^=0.00029, F(1,19)=0.05492, p=0.08172; metestrus: r^2^=0.09921, F(1,19)=2.093, p=0.1643), suggesting that this neuronal population is required for the body mass-dependent modulation of food intake during these stages.

**Figure 3:**
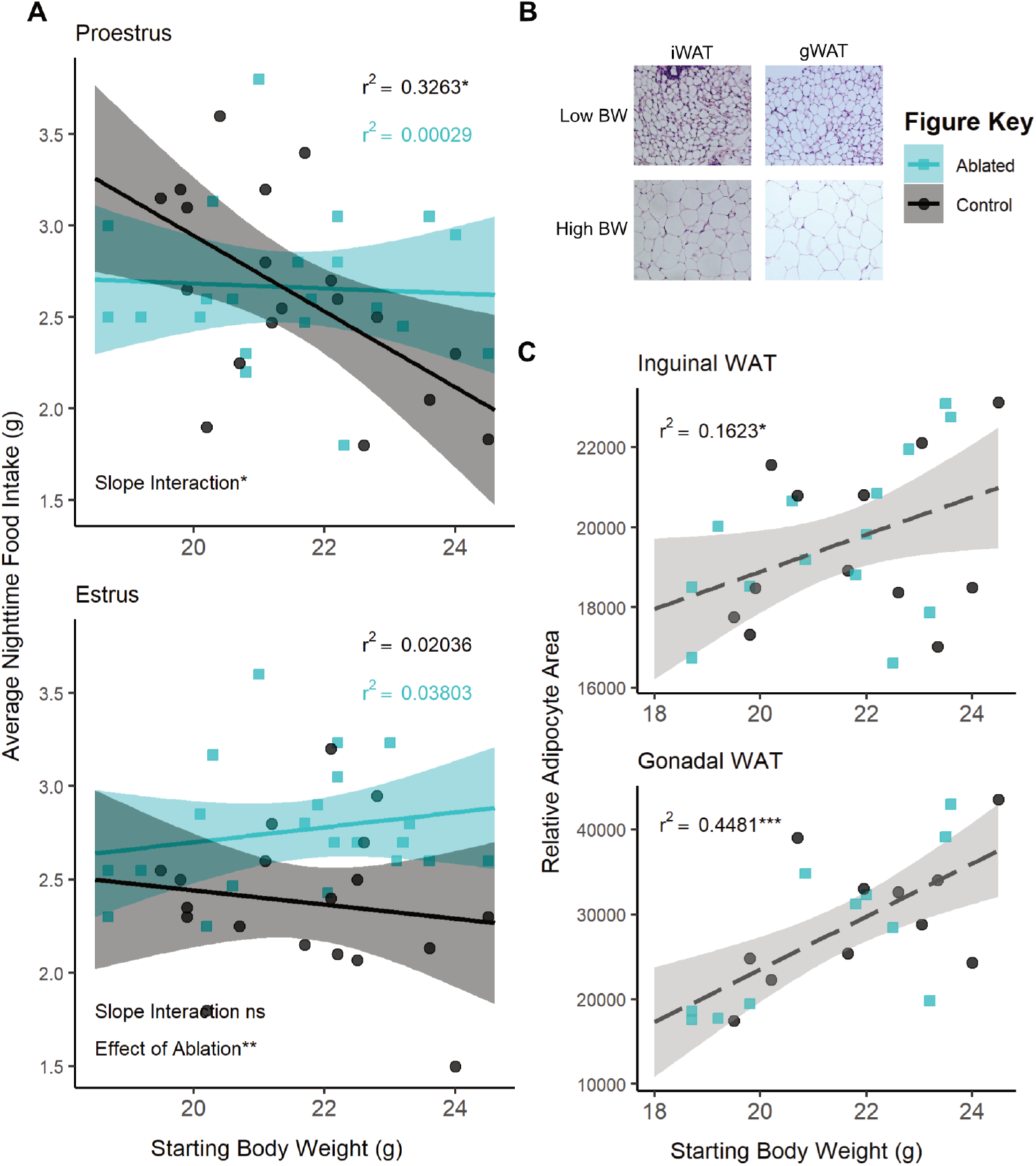
The effect of TN^SST^ neuron ablation in proestrus depends on body mass or adiposity. (A) Regression analysis within proestrus (top panel) and estrous (bottom panel) across all ovary-intact animals reveals an interaction between body mass and nightly food intake in females. Significant negative correlations in wildtype animals are seen in the high estradiol phase of proestrus but not in caspase ablated females. Control n=19, Ablated n=21 across stages. (B) Representative brightfield images of inguinal and gonadal white adipose tissue (iWAT & gWAT, respectively) in high and low body weight (BW) females. (C) Terminal iWAT (left) and gWAT (right) adipocyte size both positively correlate with starting body weight regardless of TN^SST^ neuron ablation (Interaction of slopes: iWAT F(1,22)=0.2337 p=0.6336, gWAT F(1,18)=0.4552 p=0.5085; difference in y-intercept: iWAT F(1,22)=0.1417 p=0.7103, gWAT F(1,18)=0.000 p=9971). Linear regression lines represent compiled data across ablation status. iWAT: Control n=12, Ablated n=14; gWAT: Control n=11, Ablated n=11. Mean ± 95% CI, *p<0.05, **p<0.01

Given the known influence of fat mass on feeding, adiposity was examined post-mortem. Post-mortem adipocyte size of subcutaneous inguinal and visceral perigonadal white adipose tissue (iWAT and gWAT, respectively) positively correlated with the body mass determined at the onset of feeding assays (iWAT: F(1,22)=4.3346, p=0.04919; gWAT: F(1,18)=14.9872, p=0.001119; Figure 3B&C) regardless of TN^SST^ neuron ablation (ANCOVA revealed no significant interaction of slopes or difference in y-intercept, see Table 2). Body mass accounted for a larger percentage of variation in visceral gWAT adipocyte size (r^2^=0.4481; F(1,20)=16.24, p=0.0006557) than it did for subcutaneous iWAT (r^2^=0.1623; F(1,24)=4.649, p=0.0413), suggesting a possible increased contribution of visceral adiposity to the effect of TN^SST^ neuron ablation.

### TN^SST^ neurons are sensitive to estrogens and adiposity signals

To test if TN^SST^ neurons can respond to estrogens and/or signals from white adipose tissue, we profiled the transcriptome of fluorescently labeled TN^SST^ neurons using flow cytometry followed by bulk RNA sequencing (Flow-Seq; Figure 4A). Transcriptomic analysis of isolated TN^SST^ neurons uncovered numerous differentially expressed genes between females and males, with the gene for estrogen receptor alpha, *Esr1*, being more highly expressed in females (Wχ^2^=6.736, adj p=2.356 x 10^−8^; Figure 4B). However, we were unable to detect ERα immunoreactivity in the TN using standard antibodies (Millipore Sigma 06-935). Instead, we confirmed expression from the *Esr1* locus through the injection of a Cre-dependent tdTomato reporter into the TN of female and male *Esr1-Cre* mice. Subsequent colocalization of tdTomato with *Sst* via *in situ* hybridization revealed co-expression of *Esr1* in approximately 10% *Sst*-expressing cells across sexes (Figure 4C&D), confirming that at least a subset of TN^SST^ neurons is sensitive to estrogens.

**Figure 4:**
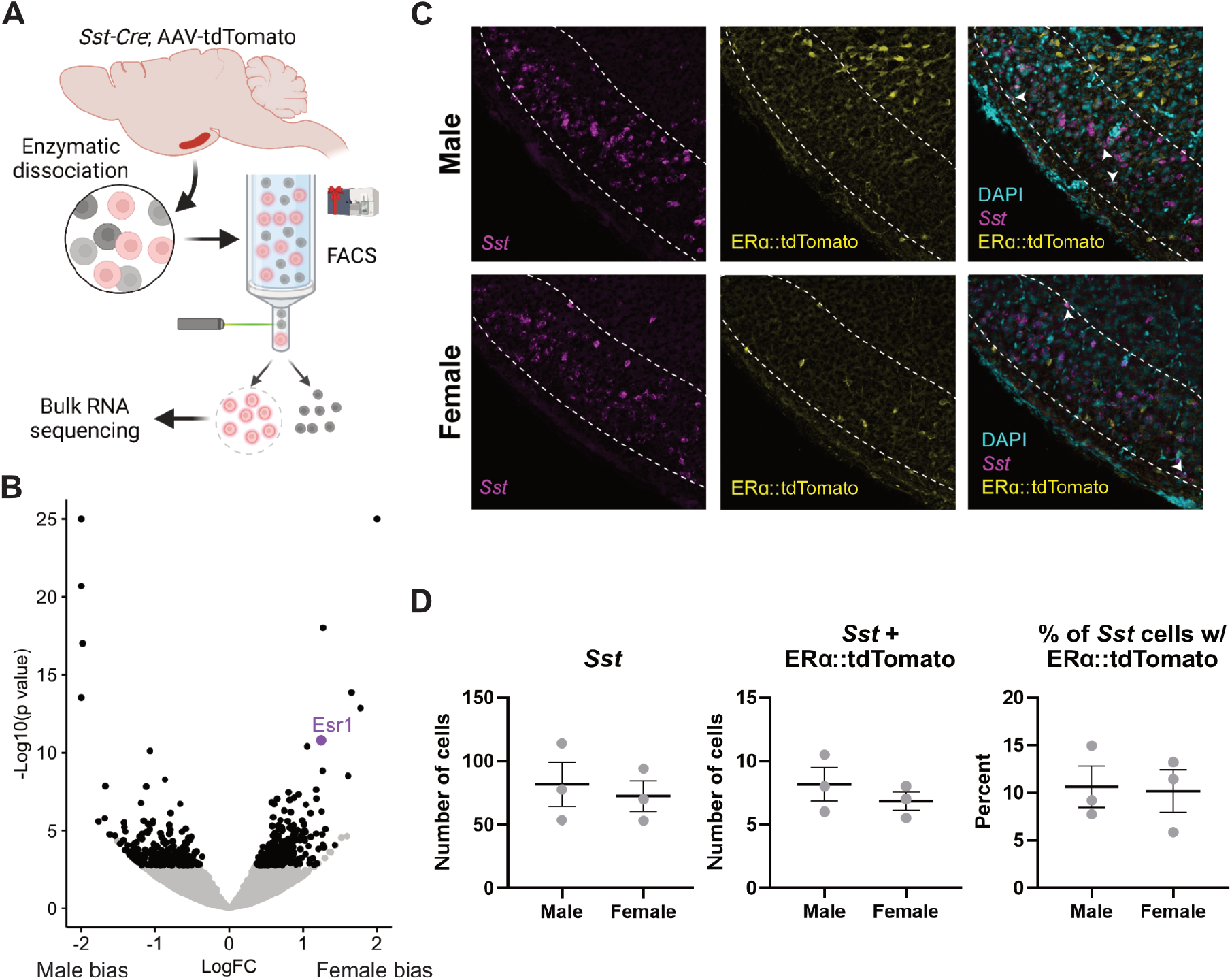
A subset of TN^SST^ neurons is sensitive to estrogens. (A) Schematic of FlowSeq analysis (also applies to Figure 5 genetic input). AAV-flex-tdTomato was stereotaxically injected into the tuberal nucleus (TN) of *Sst-Cre* mice. Lateral hypothalamic region was grossly dissected and enzymatically dissociated before segregating *Sst^+^* red neurons via flow cytometry. Resultant cells were sent for Bulk RNA sequencing. Created using BioRender.com. (B) Numerous genes are more differentially expressed between females and males (black dots), including canonical Y-associated genes and *Esr1* (purple dot). (C) Representative fluorescent images from *Esr1-Cre* mice showing colocalization of *Sst* (magenta), *Esr1∷tdTomato* (yellow), and DAPI nuclear stain (cyan). White arrows indicate *Sst-Esr1* double labeled cells. (D) Quantification of colocalization confirms *Esr1*/*Sst* co-expression but reveals no sex difference. Mean ± SEM.

To determine if TN^SST^ neurons communicate with adipose tissue or *vice versa*, we used a co-correlation analysis method based on genetic variation. High expressing genes from TN^SST^ neurons (based on counts > glial fibrillary acidic protein, GFAP, expression) were used as the “target” pathways, where human orthologues within the GTEx database (Lonsdale et al., 2013) and subjected to cross-tissue genetic co-correlational analyses (Seldin et al., 2018; Velez et al., 2022) (Figure 5A). As previous data indicated that TN^SST^ neuron responsivity to metabolic cues might be localized to periods of higher circulating estradiol (Figure 2D) and TN^SST^ may be able to directly sense circulating estradiol levels (Figure 4), individuals in the GTEx database were binned into groups with either “high” or “low” circulating estradiol levels via weighted aggregation of pan-tissue z-scores corresponding to estrogen responsive gene expression (Sup Fig 3). Binning individuals into groups with indicators of “low” or “high” estrogen signaling revealed weakly associated groups as per biweight midcorrelation (Langfelder and Horvath, 2008); bicor coefficient = −0.32, p = 0.0023), suggesting that strong cross-tissue interactions differed depending on estrogen signaling status. Next, genetic co-correlation analyses from adipose (subcutaneous & omental), skeletal muscle, stomach, and small intestine to hypothalamic highly expressed genes were conducted. A lack of significant co-correlations with small intestine resulted in this tissue being omitted from the rest of analyses. Given that subsets of strong cross-tissue correlations remained for the other tissues to highly-expressed hypothalamic DEG orthologues, relevant pathways which might contribute to signaling were examined accordingly (Figure 5). Significant interactions for co-correlations between estrogen signaling group and tissue were observed across tissues for all secreted proteins (Kruskal-Wallis test for interaction between estrogen category + tissue p=1.7×10^−218^; Figure 5B), known ligands (Kruskal-Wallis interaction p=9.9×10^−20^; Figure 5C), peptide hormones (Kruskal-Wallis interaction p=1.8×10^−13^, data not shown), and feeding behavior pathways (Kruskal-Wallis interaction p=0.0023; Figure 5D). In addition, several pathways showed specificity in strength of co-correlations from one tissue to another. For example, individuals in the higher inferred estrogen signaling group exhibited higher co-correlations between TN^SST^ and adipose within secreted proteins (p<2.2×10^−16^), ligand (p=0.025), and feeding behavior pathways (p=0.042) as compared to individuals in the low estrogen signaling group. Interestingly, this relationship was reversed for all secreted proteins in skeletal muscle, with individuals in the low estrogen signaling group exhibiting higher co-correlations (p=4.2×10^−12^). These results indicate increased communication between adipose and TN^SST^ neurons during periods of high estrogen signaling, and a switch to skeletal muscle communication when estrogen signaling is low. Across all gene sets, individuals with inferred low estrogen signaling exhibited higher co-correlations with stomach as compared to individuals with high estrogen signaling (all secreted proteins: p<2.2×10^−16^; ligands: p=1.2×10^−5^; peptide hormones: p=1.8×10^−4^; and feeding behavior: p=0.029). Together, these human genetic co-correlation data indicate that TN^SST^ neurons modulate feeding pathways through preferential communication with adipose when estrogen signaling is high and stomach hormones when estrogen signaling is decreased. These observations are in line with known responsivity of TN^SST^ neurons to stomach peptide hormone and known regulator of feeding ghrelin (Luo et al., 2018), but indicate that this communication pathway may be more salient when estradiol levels or estrogen signaling is low. Further, these analyses of human data may suggest that a similar a similar integration of metabolic cues alongside reproductive hormones in humans as well as mice.

**Figure 5:**
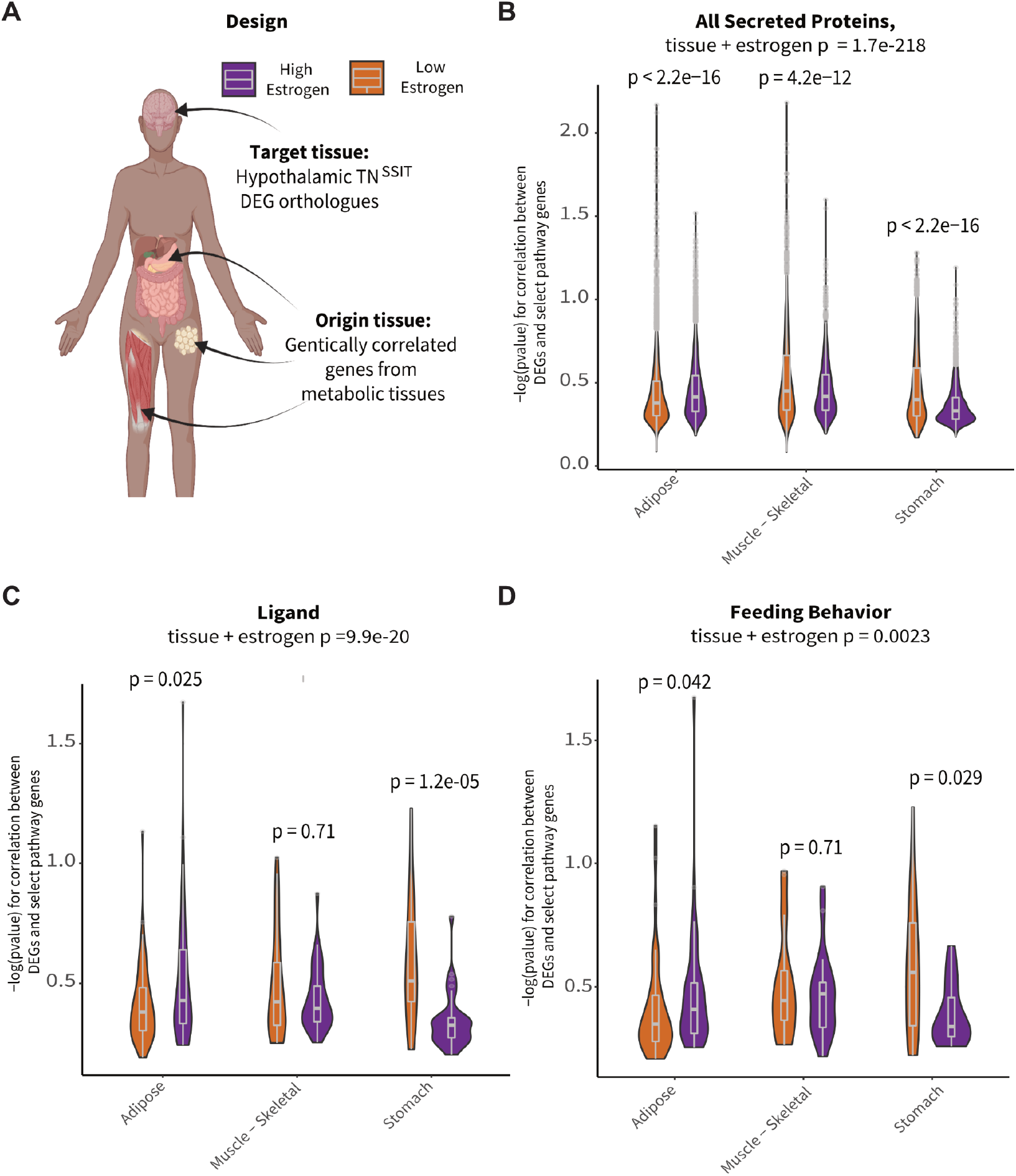
TN^SST^ neurons display increased hormonal pathway co-correlations with white adipose tissue in individuals with inferred high estrogen signaling. (A) Schematic overview of co-correlation analysis. High expressing TN^SST^ genes from mouse Flow-Seq experiments were co-correlated across various peripheral metabolic tissues across high and low estradiol groups identified in the GTEx database. Created with BioRender.com. (B) Inferred estradiol levels affected co-correlation pathways relevant to all secreted proteins. High estradiol individuals showed increased co-correlations in adipose tissue whereas those with lower estradiol showed increases in skeletal muscle and stomach communication. (C) Similar trends in adipose and stomach co-correlations across estradiol groupings were seen for ligand pathways. (D) Co-correlations across tissues showed differential impact on genes associated with feeding pathways. Individuals with higher estradiol showed increased co-correlations within these pathways with adipose tissue and decreased co-correlations with stomach as compared to those with lower estradiol levels.

To test the causal, directional relationship between fat and TN^SST^ neurons in the modulation of food intake, caspase ablation studies were repeated in combination with fat transplantation. Approximately 1.5 weeks following transplantation of ~0.8 g subcutaneous fat (Figure 6A), recipient mice exhibited significantly increased body mass (F(1,25)= 28.3184, p<0.0001; Figure 6B), raw fat mass (F(1,25)= 34.2342, p<0.0001; Figure 6B), and percent fat mass (F(1,25)=30.0008, p<0.0001; Figure 6C) and no interaction with TN^SST^ neuronal ablation in any case. Thus, fat transplantation increased adiposity similarly across neuronal ablation groups.

**Figure 6:**
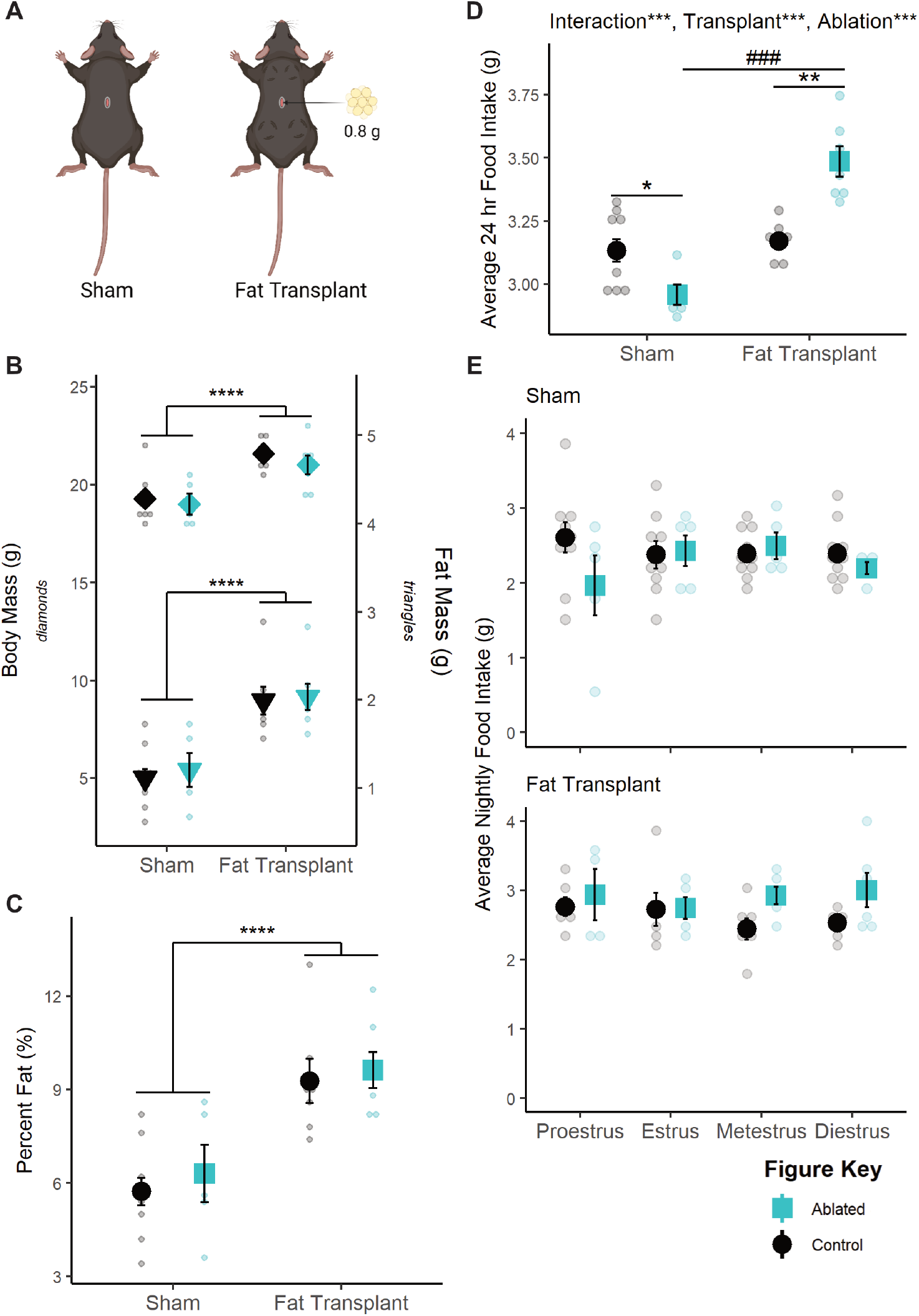
Fat transplantation modulates the effect of TN^SST^ neuron ablation. (A) Representative schematic of fat transplants. Four deposits of 0.2 g each were placed in the dorsal subcutaneous region. Created with BioRender.com. (B) Fat transplant increases raw body mass (left axis, p<0.0001) and fat mass (right axis, p<0.001) regardless of TN^SST^ neuronal ablation. (C) This translates to an overall increase in adiposity (p<0.0001). (D) Fat transplantation reverses the effect of TN^SST^ ablation, significantly increasing daily food intake compared to non-ablated controls. (E) This effect of fat transplant (top panel) seems to be due to a lack of effect during proestrus, though data were underpowered to detect the effect of estrous stage in sham controls (bottom panel). Mean ± SEM; within adiposity group: *p<0.05, **p<0.01; ***p<0.001, ****p<0.0001; between adiposity group: ^###^p<0.001. Sham: Control n=10, Ablated n=5; Transplant: Control n=7, Ablated n=7. Sham: Proestrus Control n=10 & Ablated n=5, Estrus Control n=9 & Ablated n=5, Metestrus Control n=10 & Ablated n=5, Diestrus Control n=10 & Ablated n=4; Fat Transplant: Proestrus Control n=6 & Ablated n=5, Estrus Control n=6 & Ablated n=6, Metestrus Control n=6 & Ablated n=6, Diestrus Control n=6 & Ablated n=6.

Fat transplant also increased food intake in general (F(1,25)= 52.524, p<0.0001), and the effect of TN^SST^ neuronal ablation was affected by fat transplant (F(1,25)=26.660, p<0.0001; Figure 6D). *Post-hoc* t-tests revealed that TN^SST^ neuronal ablation significantly decreased food intake in sham transplant animals (t(11.646)=−2.917, p=0.01327) but significantly increased food intake in animals receiving fat transplant (t(4.8.2536)=4.8427, p=0.001175), similar to the previous relationship with body mass in proestrus (Figure 3A). However, we were unable to detect a significant interaction with fat transplantation and ablation status over the estrous cycle (Figure 6E), possibly due to high variability in nighttime food intake in sham-transplanted mice. Together, these findings indicate that fat transplantation masks the proestrus-specific effect of TN^SST^ neuron ablation and reveal a role for fat mass in modulating the function of TN^SST^ neurons within the feeding circuit.

## Discussion

These data suggest that TN^SST^ neurons are a locus in the brain that mediates metabolic and reproductive tradeoffs. While activation of TN^SST^ neurons increases food intake across sexes, permanent inactivation by ablation during adulthood results in decreased food intake only in females during the proestrus phase. This effect depends on body mass, as this effect is apparent only in lighter animals. In wildtype mice, body mass inversely correlates with food intake on the night of proestrus, but TN^SST^ neuron ablation uncouples this effect. Further analysis reveals that white adipose tissue abundance is a significant contributing factor to this effect. Not only does post-mortem adipocyte size correlate with body mass in neuron ablation experiments, but fat transplantation studies confirm that TN^SST^ neuron ablation only decreases food intake in lean animals compared to their fat transplanted counterparts. This interaction between cycling adipokines and gonadal hormones may be mediated by the direct effects of these circulating molecules on TN^SST^ neurons, as these cells show some estrogen sensitivity via co-expression analyses. Furthermore, co-correlations between the hypothalamus and adipose tissue in humans and fat transplantation experiments in mice point to the importance of secreted proteins and ligands, suggesting TN^SST^ may detect and respond to adipokines. Future studies are needed to confirm and dissect the mechanisms of these cellular effects.

What adipokine factor is possibly being detected by the TN^SST^ remains to be determined. Leptin positively correlates with overall adiposity (Fontana and Della Torre, 2016), and it has been known to play a crucial role in reproductive responsiveness to metabolic condition, namely as the permissive signal required for pubertal onset (Chehab et al., 1996; Cheung et al., 1997). Adiponectin, an adipokine that negatively correlates with visceral fat mass in mammals (Fontana and Della Torre, 2016), has long been shown to downregulate reproduction through direct impacts on the hypothalamus (Rodriguez-Pacheco et al., 2007). Resistin is correlated with higher overall adiposity (Yang et al., 2012), exhibits numerous interactions with the hypothalamic-pituitary-gonadal axis (Mathew et al., 2018; Nogueiras et al., 2003; Tsatsanis et al., 2015), and is down-regulated during the fertile periods of the mouse estrous cycle (Gui et al., 2004).

Regardless of adipokine contributor, this trade-off paradigm provides a plausible explanation for the varied effects of estradiol on food intake in mice. While endogenous fluctuations and experimental manipulations of estradiol consistently reveal that estrogens decrease food intake in rats and guinea pigs (Asarian and Geary, 2013, 2002; Clegg et al., 2007; Eckel, 2011), the mouse literature is less definitive (Eckel, 2011; Geary et al., 2001; Naaz et al., 2002; Petersen, 1976; Witte et al., 2010). Instead, the more consistent phenotype in mice is a decrease in energy expenditure following estradiol depletion (Correa et al., 2015; Musatov et al., 2007; Xu et al., 2011). In light of this study, it is possible that the effects of estradiol on feeding across mouse studies, as observed by either endogenous estrous cycle fluctuations or ovariectomy manipulation, could be confounded by body mass and adiposity. Thus, factors like age at time of experiment, differences in fat distributions between species or strains, diet, or ovariectomy and time from ovariectomy to estradiol replacement might present confounds based on changes to fat and/or lean mass.

How circulating estrogen levels contribute to this circuit also requires further investigation. Our human GTEx analyses shows that co-correlations between TN^SST^ genes and that in peripheral tissue shifts from predominantly adipose-based to skeletal muscle- and stomach-based depending on evident estrogen signaling (Figure 5). This suggests that higher estrogen levels may increase communication between TN^SST^ neurons and white adipocyte depots, particularly as it relates to regulation of feeding behavior (Figure 5D). While this could be due to the actions of circulating estrogens on white adipose tissue itself (reviewed in (Hevener et al., 2015; Palmer and Clegg, 2015)), it is also possible that estrogens directly act on TN^SST^ neurons to increase their sensitivity and/or responsivity to adipokines. Indeed, TN^SST^ neurons exhibit estrogen sensitivity (Figure 4), though future studies would be needed to test for a possible direct effect.

Alternatively, fluctuating hormone levels might be detected elsewhere in the brain and impact TN^SST^ neuronal modulation of feeding through integration at the circuit level. TN^SST^ neurons project to many estrogen-sensitive nodes or nodes receiving direct input from estrogen-responsive regions, including the bed nucleus of the stria terminalis, parabrachial nucleus, and central amygdala (Luo et al., 2018). This circuit-wide integration of estradiol is a known mechanism of action for the gonadal hormone, with estrogens acting on many circuit nodes to coordinate behavioral output in a variety of cases, including reward/addiction (Becker and Chartoff, 2019) and thermoregulation (Zhang et al., 2021). It is therefore probable that the effects of estradiol on feeding function similarly, as the anorexigenic effects of estradiol have been localized to numerous feeding nodes such as the hypothalamic arcuate nucleus (Roepke et al., 2010, 2007; Santollo et al., 2011; Todd L Stincic et al., 2018; Todd L. Stincic et al., 2018) and the nucleus of the solitary tract of the brainstem (Asarian and Geary, 2006; Maske et al., 2017).

In all, this study adds to the growing literature interrogating the contributions of TN^SST^ neurons to feeding behavior. Central SST (originally named growth hormone inhibiting hormone, GHIH, in the central nervous system, (Painson and Tannenbaum, 1991) had long been known to affect food intake through somatostatin receptor 2 (SSTR2; (Beranek et al., 1999; Campbell et al., 2017; Danguir, 1988; Lin et al., 1989; Andreas Stengel et al., 2010b, 2010a; A Stengel et al., 2010; Stengel et al., 2015, 2013, 2011; Tachibana et al., 2009). This effect was seemingly localized to the tuberal nucleus, when TN^SST^ neurons were found to integrate into the melanocortin feeding system, though the effect of these neurons on feeding was attributed to γ-aminobutyric acid (GABA) release as opposed to direct SST effects (Luo et al., 2018). Subsequently, TN^SST^ neurons were also found to contribute to food context learning in males^1^ (Mohammad et al., 2021), indicating that the TN may straddle homeostatic and hedonic feeding mechanisms (Massa and Correa, 2020). This paper adds to this growing literature by not only delineating an apparent sex difference but also context dependence in TN^SST^ neuronal modulation of food intake.

We further speculate that TN^SST^ neurons serve as a nexus of integration and a mediator of reproductive and metabolic tradeoffs within the feeding circuit. In cycling rodents, fertile periods during the estrous cycle are accompanied by alterations to metabolic output, including a decrease in food intake (Asarian and Geary, 2013, 2002; Brobeck et al., 1947; Eckel, 2011), increase in locomotion (Brobeck et al., 1947; Kent et al., 1991; Sanchez-Alavez et al., 2011; Steiner et al., 1982), and increased core body temperature (Kent et al., 1991; Sanchez-Alavez et al., 2011). These changes are hypothesized to suppress energy intake and promote active mate-seeking behavior and sexual receptivity. This study identifies TN^SST^ neurons as possible mediators of such a trade-off, actively promoting energy intake during fertile periods when metabolic reserves may be insufficient to support reproduction.

## Materials & Methods

### Mice

Female (defined as having small anogenital distance at weaning and presence of ovaries at time of death) and male (defined as having large anogenital distance at weaning and presence of testes postmortem) mice expressing the *Sst-Cre* driver transgene (JAX stock no. 013044, *Sst^tm2.1(cre)Zjh^*/J) were maintained on a C57BL/6J genetic background. Heterozygotes (*Sst-Cre/+*) and/or wildtype littermates (*+/+*) were used for all studies. Genotypes were determined as per JAX protocol 28317. Female and male mice expressing the *Esr1-Cre* driver transgene (JAX stock no. 017911, B6N.129S6(Cg)-*Esr1^tm1.1(cre)And^/J*) were maintained on a C57BL/6J genetic background. Heterozygotes (*Esr1-Cre/+*) were used for colocalization studies. Genotypes were determined as per primers from JAX protocol 27213. Experiments were performed on cycling females and gonadally-intact males unless otherwise stated. Mice were maintained on a 12:12 light cycle, with *ad libitum* access to food and water (unless otherwise specified), under controlled humidity conditions, and in single-housed cages with non-caloric paper bedding to ensure accurate food intake assessment. All studies were carried out in accordance with the recommendations in the Guide for the Care and Use of Laboratory Animals of the National Institutes of Health. UCLA is AALAS accredited, and the UCLA Institutional Animal Care and Use Committee (IACUC) approved all animal procedures.

### Estrous cycle staging

Vaginal lavages were performed on females daily, between ZT 0 and ZT 4, using 30 μL of standard phosphate buffered saline (PBS). Samples were deposited onto slides and allowed to dry prior to staining. Males were subjected to similar handling during this time to ensure roughly equivalent handling stress. Giemsa staining was carried out to visualize cellular composition of the vaginal cavity. Stock Giemsa stain was prepared at least one week in advance of use. An 18.5% solution of Giemsa powder (Fisher G146-10) in glycerin was heated to 60°C and cooled before diluting 9:14 with 100% methanol. Stock was diluted 1:30 in PBS before use, shaking vigorously before stain. Slides were incubated for one hour at room temperature. Prior to staining, slides were briefly fixed in 100% methanol. Staging was assessed via light microscopy as in (Cora et al., 2015), and stages were assigned using the behavioral method (Becker et al., 2005), with morning samples indicating the prior night’s estrous stage. This staging method was confirmed by core body temperature waveform alignment (Sanchez-Alavez et al., 2011).

### Surgical Procedures

Mice received analgesics (0.074 mg/kg buprenorphine two times daily, 7.11 mg/kg carprofen one time daily) on the day of and one day post-surgery. Mice were anaesthetized with 3% isoflurane and maintained within a range of 1.25-2.5%. AAVs were bilaterally injected into the TN of adult mice (coordinates relative to Bregma: A-P −1.65 mm, lateral ±0.75, D-V −5.45; scaled when Bregma-Lambda distance was not equivalent to 4.2 mm) at a rate of 5 nL/s using a glass-pulled needle. See Table 1 for titers and injection volumes. Controls consisted of both wildtype animals injected with the experimental virus (virus controls) and Cre positive animals injected with cell-filling GFP (genotype controls). Ovariectomy surgeries included complete removal of gonads from adult mice. Gonadectomies occurred immediately prior to stereotaxic viral injections within the same surgical period. In telemetry experiments, G2 eMitters (Starr Life Sciences) were implanted intraperitoneally on the same day as viral injection. Experiments were conducted following at least two weeks recovery from surgical proceedings.

**Table 1:**
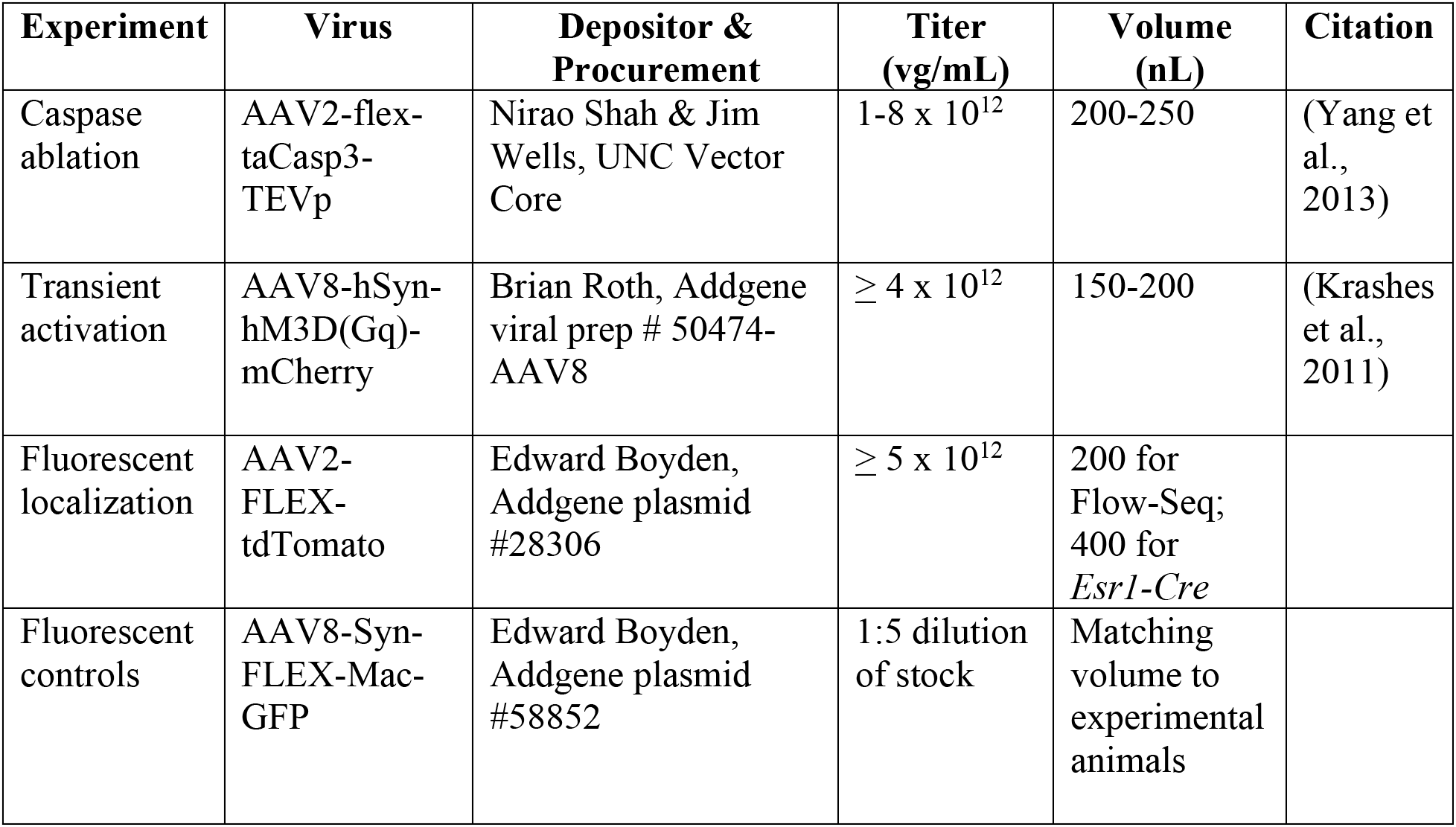
List of viral vectors used.

### Caspase ablation experiments

Gross movement and core body temperature were passively measured every other week for eight weeks using VitalView software (Starr Life Sciences). Body weight was measured every week. Food assay was performed when mice were not on telemetry pads. At ZT 0.5 on the start day of the experiment, 2/3 of the non-caloric paper bedding was removed. A pre-measured amount of food was delivered, and mouse body weight measured. Food in hopper was weighed at ZT 0.5 and ZT 11.5 every day until experiment conclusion. After 96 hours, food and all bedding were collected to account for food spillage. For some experiments, 4-5 hour fasted glucose tolerance tests were performed prior to sacrifice. In ovariectomy experiments, two food assays were performed back-to-back, non-fasted resting glucose levels were collected, body composition was measured via NMR, and indirect calorimetry was performed in Oxymax metabolic chambers (Columnbus Instruments) at room temperature. Upon experiment completion, all brains were collected using RNase-free conditions. Inguinal white adipose tissue (iWAT) and gonadal white adipose tissue (gWAT) were collected for histology analyses.

### Transient activation food intake assay

Clozapine-n-oxide (CNO; MilliporeSigma #0832) was used to activate TN^SST^ neurons in *Sst-Cre* animals expressing hM3Dq-mCherry. Stock solution of 20 mg/mL in DMSO was stored at −20°C and diluted to a working solution of 0.03 mg/mL in sterile saline also stored at −20°C. Saline control (0.15% DMSO) or CNO (10 μL/g body weight, dose of 0.3 mg/kg) working solution were administered IP in a counterbalanced design. Experiments were completed in duplicate replicate trials. Mice were transferred to experimental room at least 15 minutes prior to experimentation. Experiments were begun between ZT 2-3 and terminated between ZT 6-7. Following injection, food intake was measured at 0.5, 1, 2, and 4 hr. Vaginal lavage was performed on female mice after experiment conclusion to prevent stress interference with food intake. All mice were injected with CNO 90 minutes prior to sacrifice to enable neuronal activation validation via cFOS immunohistochemistry.

### Fat Transplantations

Donor fat was taken from various visceral (i.e., periuterine perigonadal, retroperitoneal, and omental) depots of wildtype female C57BL/6J mice and implanted into female mice recently stereotaxically injected under standard surgical conditions. Four depots of 0.15-0.25g were placed subcutaneously on the dorsal surface through a single incision mid-back, for a final transplantation total of 0.6-0.9g of white adipose. Fat for each depot was divided into at least three individual pieces to promote vascularization. The visceral-to-subcutaneous paradigm was used due to the deleterious metabolic effects of this graft (Tran et al., 2008). Food intake was assayed 2-3 weeks following transplantation to allow for sufficient angiogenesis (Gavrilova et al., 2000) and graft stabilization without endogenous fat depot compensation (Rooks et al., 2004). Upon sacrifice, grafts were examined to confirm tissue was not necrotic.

### Histology

#### In situ hybridization (ISH) and immunostaining (IHC)

RNA probe generation was accomplished as in (van Veen & Kammel et al., 2020). Briefly, *Sst* sense and antisense probes were transcribed using a DIG or FITC RNA labeling kit (Roche) and purified with RNA Clean & Concentrator (Zymo Research). PCR products were amplified using Allen Brain Institute-derived reference primer sequences and cloned into pCR 2.1 TOPO (Invitrogen). Plasmid DNA was then isolated from bacterial cultures (ZymoPURE II Plasmid Midiprep kit), linearized, and purified (Zymo DNA Clean & Concentrator). Validation of caspase ablation was carried out on 35μm-thick coronal slices via chromogen ISH protocol was as per (van Veen & Kammel et al., 2020). Validation of hM3Dq targeting and activation was accomplished by visualization of native mCherry expression and IHC stain for cFOS. Briefly, slides were blocked and incubated with rabbit anti-cFOS (1:200, Synaptic Systems # 226003, RRID: 2231974) primary antibody overnight at 4°C. The next day, sections were incubated for 1 hour at room temperature with goat anti-rabbit Alexa Fluor 488 secondary (1:500, Thermo Fisher Scientific # A11034, RRID: AB_2576217) and counterstained with DAPI. For colocalization experiments, *Esr1-Cre* mice were bilaterally injected with 400 μl AAV2-flex-tdTomato into the TN coordinates. Native tdTomato fluorescence destroyed by combined ISH protocol was recovered by rabbit anti-DsRed (1:1000, Takara Bio Clontech # 632496, RRID: AB_10013483) antibody and switched to the green channel using an Alexa Fluor 488 secondary. Dual *Sst* ISH & tdTomato IHC protocol was accomplished via TSA amplification. Briefly, 35 μm sections were fixed, permeabilized with Triton X-100, and acetylated before overnight ISH probe incubation at 65°C. The next day, tissue was then washed, blocked with Blocking Reagent (MilliporeSigma 11096176001Roche) and heat inactivated sheep serum, and incubated with anti-DsRed overnight at 4°C. The final day, tissue was washed before ISH signal was developed with the TSA Plus Cyanine 5 System (Akoya Biosciences # NEL745001KT). Slides were then stripped of horseradish peroxidase and blocked with normal goat serum before incubating with goat anti-rabbit Alexa Fluor 488 (1:400) for 2 hours at room temperature.

#### Adipocyte size quantification

Inguinal and white adipose tissue was collected post-mortem and drop-fixed in 4% paraformaldehyde (PFA) for at least 18 hours. Tissue was then washed in PBS before being stored in PBS at 4°C until tissue analysis. For histological processing, tissue was placed in tissue processing cassettes and submerged in 70% ethanol before being embedded in paraffin, sectioned at 4 μM, and stained with hematoxylin & eosin (H&E) by the UCLA Translational Pathology Core Laboratory. Three regions of interest per tissue-type per mouse were imaged by light microscopy at 20x magnification. Adipocyte area was quantified using a custom pipeline in CellProfiler. Inclusion parameters were cell diameters of 100-300 pixel units and a global threshold strategy with minimum cross-entropy.

#### Colocalization analysis

*Sst* and *Esr1* co-expression was determined using CellProfiler (version 4.2.1). First, a contour was drawn around a matched section of the TN using anatomical landmarks (i.e., shape of arcuate nucleus and third ventricle). For each hemisphere, DAPI-stained nuclei were detected and intensity thresholding was used to determine *Sst+* cells. Incorrectly labeled cells were manually erased or added. *Sst+* cells were then filtered based on *Esr1∷tdTomato* signal intensity. Counts were made of total *Sst+* cells, as well as *Sst*+/*Esr1*+ and SST+/ *Esr1*- cells. The counts were averaged across the two hemispheres, and percent was calculated as ([*Sst*+/*Esr1*+] / Total SST) × 100.

### Bioinformatics Analysis

*Sst-Cre* female and male mice were bilaterally stereotaxically injected with AAV expressing Cre-dependent tdTomato (See Table 1). Following at least two weeks for viral expression, animals were sacrificed and TN was microdissected under fluorescent illumination. Dissected TN was dissociated using a papain-based enzymatic process (Worthington Biochemical) and then TN^SST^ neurons were enriched and collected via flow cytometry. Cells were sorted from debris and doublets were excluded by gating on forward-scatter and side-scatter profiles. Live nucleated cells were then selected by DAPI-negative (live) and DRAQ5-positive (nucleated) staining. Finally, tdTomato-positive cells were selected based on relatively high levels of red fluorescence (as in van Veen & Kammel et al., 2020). RNA was isolated from 500-2500 cells by RNeasy Micro kit (Qiagen). Cells were then submitted for bulk RNA sequencing. Single-end reads (~10 million unique reads per mouse) were assembled to the mouse transcriptome (version mm10) using kallisto (version 0.46.2). Differentially expressed genes and normalized read counts were identified using DEseq2 Galaxy Version 2.11.40.6+galaxy1. Volcano plots were produced by the custom R function “deseq_volcano_plot_gs()” available through the following package: http://github.com/jevanveen/ratplots. Raw reads of the RNA sequencing data were also examined for hypothalamus-peripheral tissue co-correlations across stomach, small intestine, skeletal muscle, visceral fat, and subcutaneous fat as hypothalamic reads using the GTEx database as previously described (Seldin et al., 2018; Velez et al., 2022). In addition, estrogen-responsive genes used to infer “low” vs “high estrogen signaling were gathered from: https://www.gsea-msigdb.org/gsea/msigdb/cards/HALLMARK_ESTROGEN_RESPONSE_EARLY.html. To clarify the analysis, estrogen signaling binning per individual and subsequent cross-tissue correlations with mouse DEG orthologues, all processed datasets, scripts used to analyze, and detailed walk-through is available at: https://github.com/Leandromvelez/sex-specific-endocrine-signals.

### Statistical Analyses

All statistics were carried out in R. Sex differences were determined by interaction terms between genotype and sex (caspase ablation experiments) or genotype, treatment, and sex (chemogenetic experiments). In caspase ablation and fat transplantation experiments, animals meeting both the criteria of outlier by Cook’s distance, as well as “miss” (no hit or unilateral hit as defined by more than 5% of targeted cells still present) were excluded. For fat transplantation studies, only sham animals with <10% fat mass at the beginning of the feeding assay were included. All data were checked and transformed, if necessary, to meet normalcy criteria.

## Supporting information

Table 2 - Statistical Details

## Acknowledgements

This research was supported by the UCLA Life Sciences Division (NIH R01 AG066821), the NIH Specialized Centers of Research Excellence (SCORE) grant U54 DK120342, and the National Center for Advancing Translational Science (NCATS) under the UCLA Clinical and Translational Science Institute grant UL1TR001881. MGM was supported by an NSF GRFP (DGE-2034835), Dissertation Year Fellowship from the UCLA Graduate Division, and the UCLA Kenneth I. Shine Fellowship. SA was supported by the CARE program of the UCLA Undergraduate Research Center (NIH NIGMS IMSD GM055052; PI: Hasson). MS was supported by NIH grant DK130640. The authors also wish to thank Drs. A. Arnold, K. Wassum, and E. Hsiao for helpful feedback. MGM would also like to thank all members of the Correa lab who ever assisted with timed food assays.

---

**Supplementary Figure 1 (companion to Figure 2):**
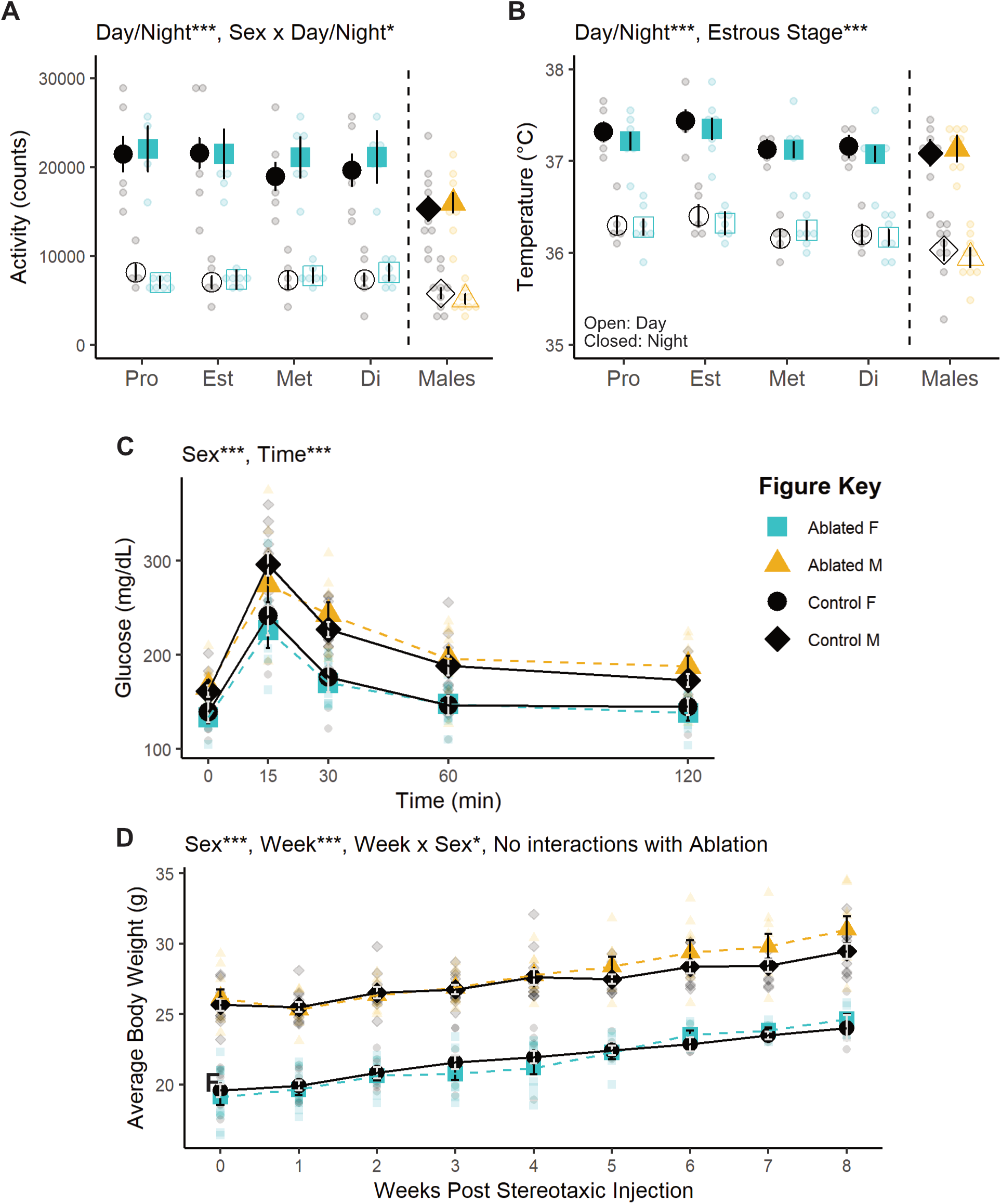
TN^SST^ neuronal ablation does not affect any other metabolic measures studied. Telemetry measures of (A) activity and (B) core body temperature are unaffected by TN^SST^ neuronal ablation. (C) Fasting glucose tolerance test is also unaffected by TN^SST^ neuron ablation. (D) Despite changes to food intake, ablation does not affect body weight over time. Mean ± SEM; *, p<0.5; ***, p<0.001. M Control n=9; M Ablated n=9; F Control n=5; F Ablated n=7. Pro: Proestrus, Est: Estrus, Met: Metestrus, Di: Diestrus.

**Supplementary Figure 2 (companion to Figure 3).**
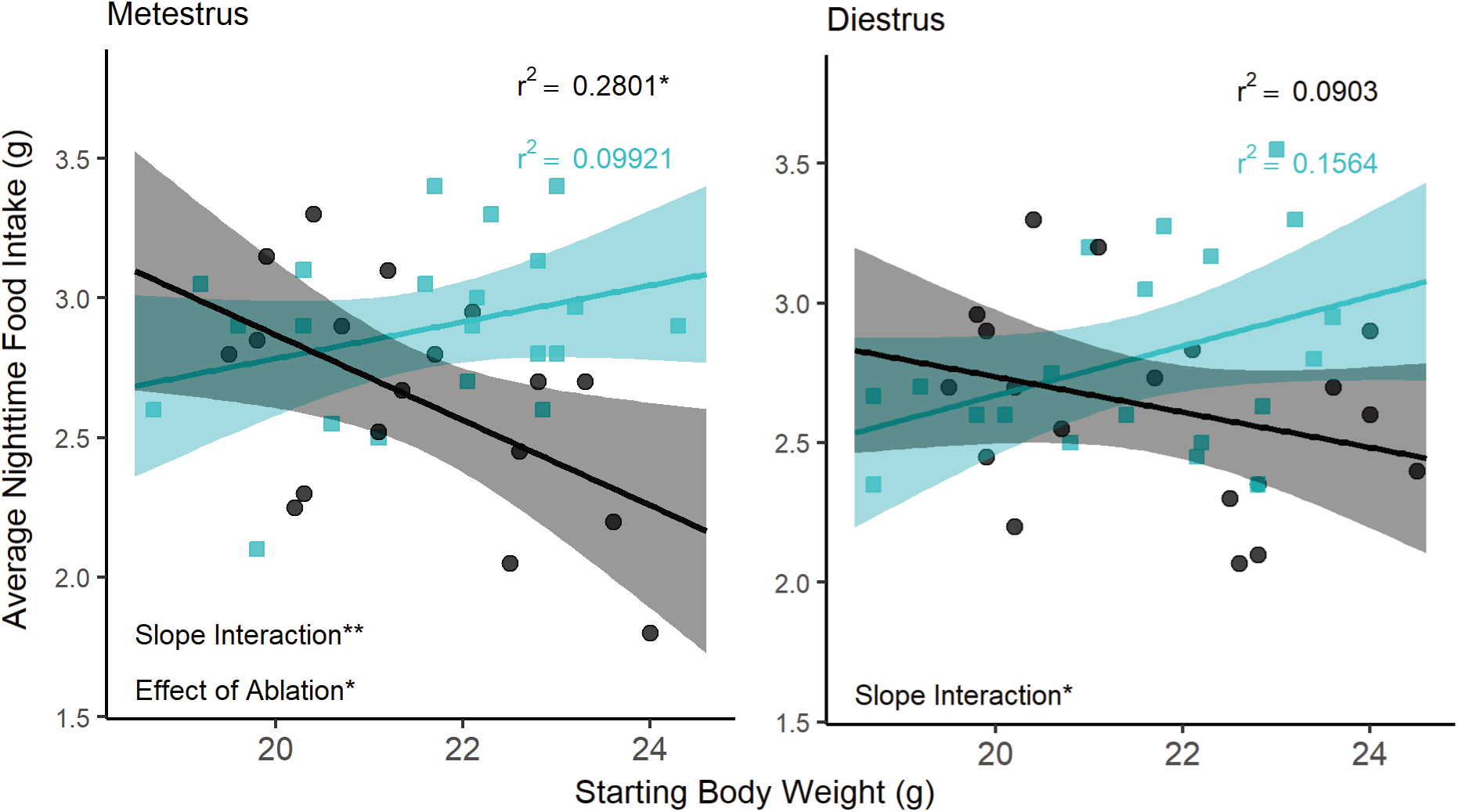
(A) Regression analysis of food intake and body mass across all ovary-intact animals in metestrus (left panel) and diestrus (right panel) reveals an interaction between body mass and nightly food intake in females. Significant negative correlations in wildtype animals (black line, round black dots) are seen in the higher estradiol phase of metestrus but not in caspase ablated females (cyan line, square cyan points). Metestrus: Control n=18, Ablated n=21; Diestrus: Control n=19, Ablated n=20.

**Supplementary Figure 3 (companion to Figure 5).**
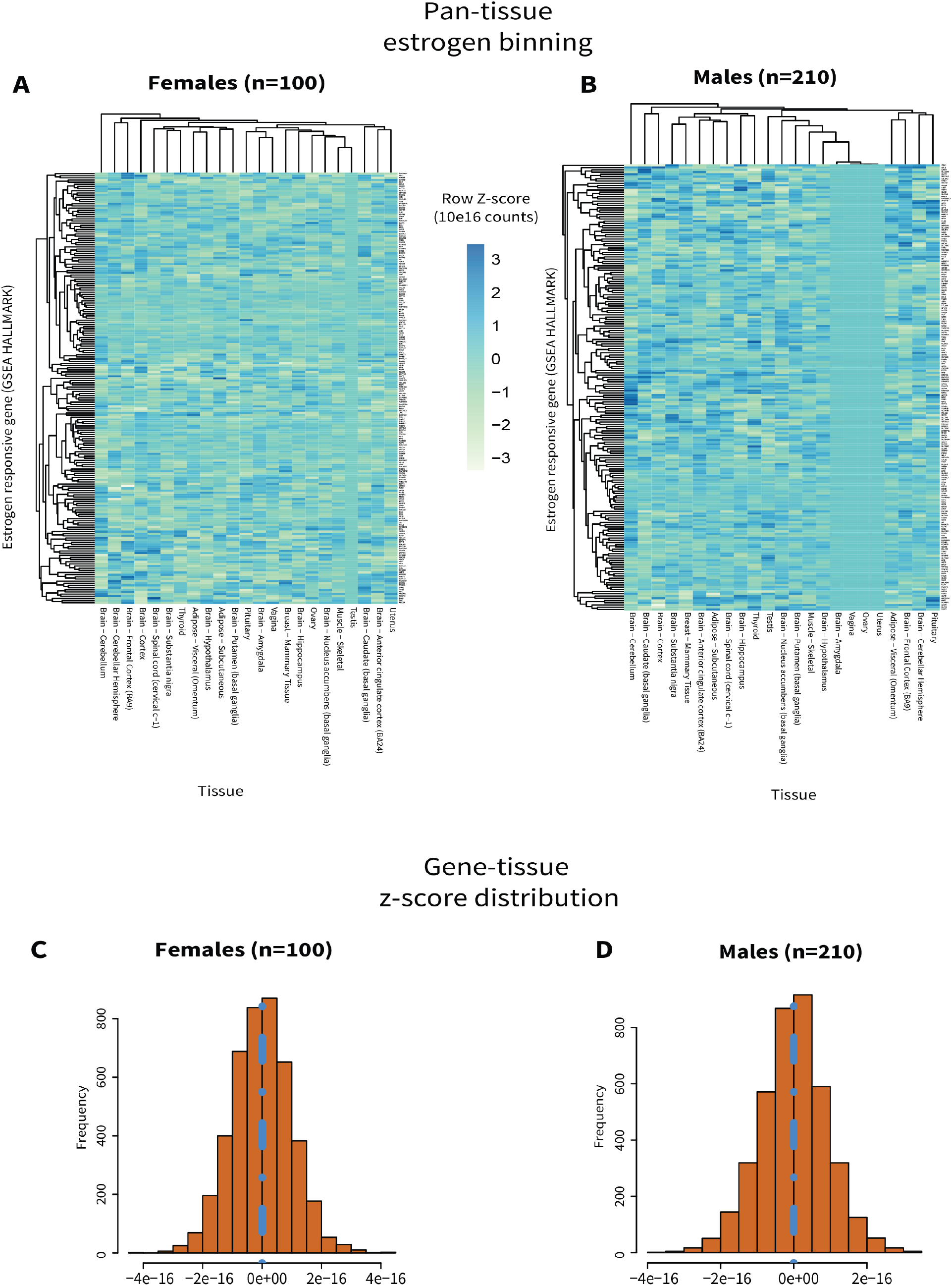
(A&B) Z-scores of estrogen signaling genes (y-axis, https://www.gsea-msigdb.org/gsea/msigdb/cards/HALLMARK_ESTROGEN_RESPONSE_EARLY.html) across indicated tissues (x-axis) in GTEx female (no Y chromosome, A) and male (Y chromosome present, B) individuals. (C&D) Based on the distributions of these scores across relevant metabolic tissues, individuals were segregated into categories of “low” (<0, left of blue line) or “high” (>0, right of blue line) estrogen signaling and used for cross-tissue genetic correlations.

## Footnotes

1 In all external papers discussed, no definitions for sex category were ever provided. In mice, we assume that sexes were defined using anogenital distance.

